# Understanding pigeon pea (*Cajanus cajan*) production conditions, stakeholders’ preferences for varietal traits and their implications for breeding programmes in India

**DOI:** 10.1101/2020.06.08.139832

**Authors:** A. Singh, I. Fromm, G. K. Jha, P. Venkatesh, H. Tewari, R. Padaria, U. Egger

## Abstract

Pigeon pea (*Cajanus cajan*) is an important pulse crop in the Indian diet and one of the most important sources of dietary protein for the population. In the context of the fourth phase of the Indo-Swiss Collaboration in Biotechnology, an assessment how farmers and consumers perceive new pigeon pea cultivars and what are their preferred traits was conducted. This investigation assessed India’s food security implications due to stagnating and low yield of pigeon pea and ascertain farmers’ preferences of pigeon pea varietal traits, production constraints and farmers’ coping strategies in diverse pigeon pea production environments. Results indicated that production constraints in the studied regions were basically similar, with majority of the farmers identifying pod borer & pod fly as the major pest, and wilt as the major disease and drought as a major production constraint. Farmers indicated the use of clean seed, high yielding varieties, inter & mixed cropping, planting density, and manure application as some of the strategies they used to cope with the production constraints. In terms of preference for new cultivars, farmers want high grain yield with drought tolerance, medium to early maturity, pod borer resistance, tolerance to wilt disease, moderate plant height and ease of threshing without compromising other preferred attributes. The analysis of processors’ preference of grain types for dal processing showed that uniform size, oval shape, orange coloured seed and most importantly moderately hard seed coat are highly desirable. Consumers lacked awareness on the varieties, but considered traits like uniform seeds and reduced cooking time traits more desirable.

## Introduction

Pulses, being a major source of protein, constitute an integral component of Indian’s dietary basket and are grown under diverse agro-climatic conditions of the country. Among pulses, pigeon pea (*Cajanus cajan)* stands second in area and production, next only to chickpea. Pigeon pea thrives in hot dry environments, its drought tolerance and ability to utilize residual moisture with low inputs during the dry season makes it important for the livelihood of smallholders in the semi-arid tropics (Jones *et al*.,2002). Globally, pigeon pea is grown on 7.02 million hectares (m ha), mainly in Asia, Latin America and Eastern and Southern Africa with an average yield of 0.97 t/ha in the year 2017 (FAOSTAT, 2017). However, pigeon pea occupies a unique place on Indian agriculture scene as the country accounts for about 71.5% of the global production, covering an area of around 5.40 m ha (occupying about 15.4 % of area under pulses) and production of 4.78 million tonnes (mt) (contributing to 20.9 % of total pulse production). Global yield largely reflects the situation in India where pigeon pea yield stagnated around 0.70 t/ha for the last several years. Since it is mainly a rainfed crop, unfavourable rainfall (delayed & erratic) leads to terminal moisture stress, hence affecting productivity. Low seed replacement rates, non-adoption of improved management practices and lack of commercial perspective for the crop also influence the productivity to a greater extent (Sameer Kumar et al., 2014). As a multiple purpose drought-tolerant crop, pigeon pea provides many benefits to resources-poor families including protein-rich grain, fuel, fodder, fencing material, improved soil fertility and control of soil erosion. In India, pigeon pea is mainly cultivated by smallholder farmers in the arid and semi-arid lands, primarily as a source of food and cash. It is mainly consumed as dhal^1^ throughout the country besides several other uses of various parts of pigeon pea plant.

Currently, pulses research attracts less public attention than research on cereals and horticultural crops. In terms of legume breeding programmes, pigeon pea lags further behind chickpea. The chickpea is amongst the most researched crop worldwide unlike pigeon pea. Considering the high importance of pigeon pea, Indian Council of Agricultural Research (ICAR) started “All India Coordinated Pigeon Pea Improvement Project” in 1965. Under this mega project, crop improvement activities were simultaneously carried out in 31 research centres in diverse agro-ecological zones.

Although several varieties showed high potential under research environment, their performances under field conditions are poorly documented owing to numerous biotic and abiotic stresses the crop faces under marginal production environment. Despite the fact that a large number of high yielding varieties have been released and India is one of the largest producers of pigeon pea, the productivity of the crop remains stagnant around 0.7 t/ha as compared to its potential yield (1.5-3.0 t/ha). The production has been hovering around 3 mt annually since the 1990s, however, it reached to 4.78 mt first time with average productivity of 0.88 t/ha during the year 2016-17* [Govt. of India, 2017]. Hence, domestic production has not been able to match demand thereby making the country a net importer of pigeon pea in recent times.

Improved crop varieties may be high yielding, but their limited adoption points that they may not be attractive to farmers unless they possess other specific traits that farmers consider important as well (Asrat *et al*., 2009). Farmers’ adoption decisions are therefore not only driven by higher yield criterion but rather on complex processes that are affected by several socio-economic and psychological factors (Willock *et al.* 1999; Traxler and Byerlee, 1993). Their varietal traits preferences are the reflections of their requirement for the product which should essentially be a part of demand-driven research agenda. Studies on farmers’ crop variety choices indicate that farmers consider crop as a bundle of multiple characteristics (Wale *et al*, 2005, Smale *et al*., 2001; Edmeades *et al*., 2008; Badstue *et al*., 2003) which includes production characteristics such as disease and pest resistance, high yielding, early maturity and adaptability to unfavourable environments (Manu–Aduening *et al*., 2005); consumption and market characteristics such as taste, seed size, colour and more. There are umpteen number of literature available explaining the production performance of pulses, the determinants of adoption of improved pulse varieties by farmers in developing countries (Dankyi and Agyekum, 2007; Owusu and Donkor, 2012) (Conley and Udry, 2002; Mather *et al*., 2003; Faturoti *et al*., 2006; Badal *et al*., 2007), but the research on preferences of farmers and other stakeholders for varietal traits have received a scanty attention. Moreover, economists investigating varietal adoption have focused on the role of farm, farmer, household, physical environment and economic factors in explaining adoption behaviour (Nkonya et al., 1997; Doss et al., 2003; Muyanga, 2009; Finger et al., 2009).

## Materials and Methods

### Sampling and Farmers survey

A multi-stage random sampling procedure was used to select the sites for study. The analysis on spatial and temporal changes in pigeon pea production clearly suggests its concentration in central and southern zones i.e. states of Maharashtra, Karnataka, Madhya Pradesh, Telangana and Gujarat. Accordingly, one state in each zone i.e. Maharashtra from Central zone and Telangana from Southern zone were selected for study, representing two different agro-ecological and socio-economic environments of pigeon pea production. In Maharashtra, two districts namely Latur (longitude: 76.733°E and latitude 18.432°N) and Parbhani (longitude: 76.641°E and latitude 19.264°N) were selected randomly (Figure 1). Similarly, in Telangana, Vikarabad (longitude: 77.904 °E and latitude 17.336 °N) and Nagarkurnool (longitude: 78.304 °E and latitude 16.485°N) districts were randomly selected for detailed farmers’ survey (Figure 1). Two blocks from each selected district and two villages from each selected block were randomly chosen to identify the sample pigeon pea growers for a detailed survey. Finally, 15 respondents from each village were selected randomly. In all, 120 farmers were surveyed from each state, thus a total of 240 respondents were selected for a detailed survey in March-May 2018, just after farmers harvested the crop. The information collected included, household compositions and characteristics, household farm assets, crop and livestock production, pigeon pea cost of cultivation & production practices, production constraints, access to information and preferences for pigeon pea varietal traits etc. The participatory farmers’ discussion helped to identify a set of key traits that would be needed to develop varieties that meet end-user needs. Additional relevant data was also collected from other stakeholders in the pigeon pea value chain such processors (dhal millers and consumers). A total of 600 consumers were interviewed in two urban centres two major metropolitan cities of Delhi and Hyderabad.

**Figure 1.**
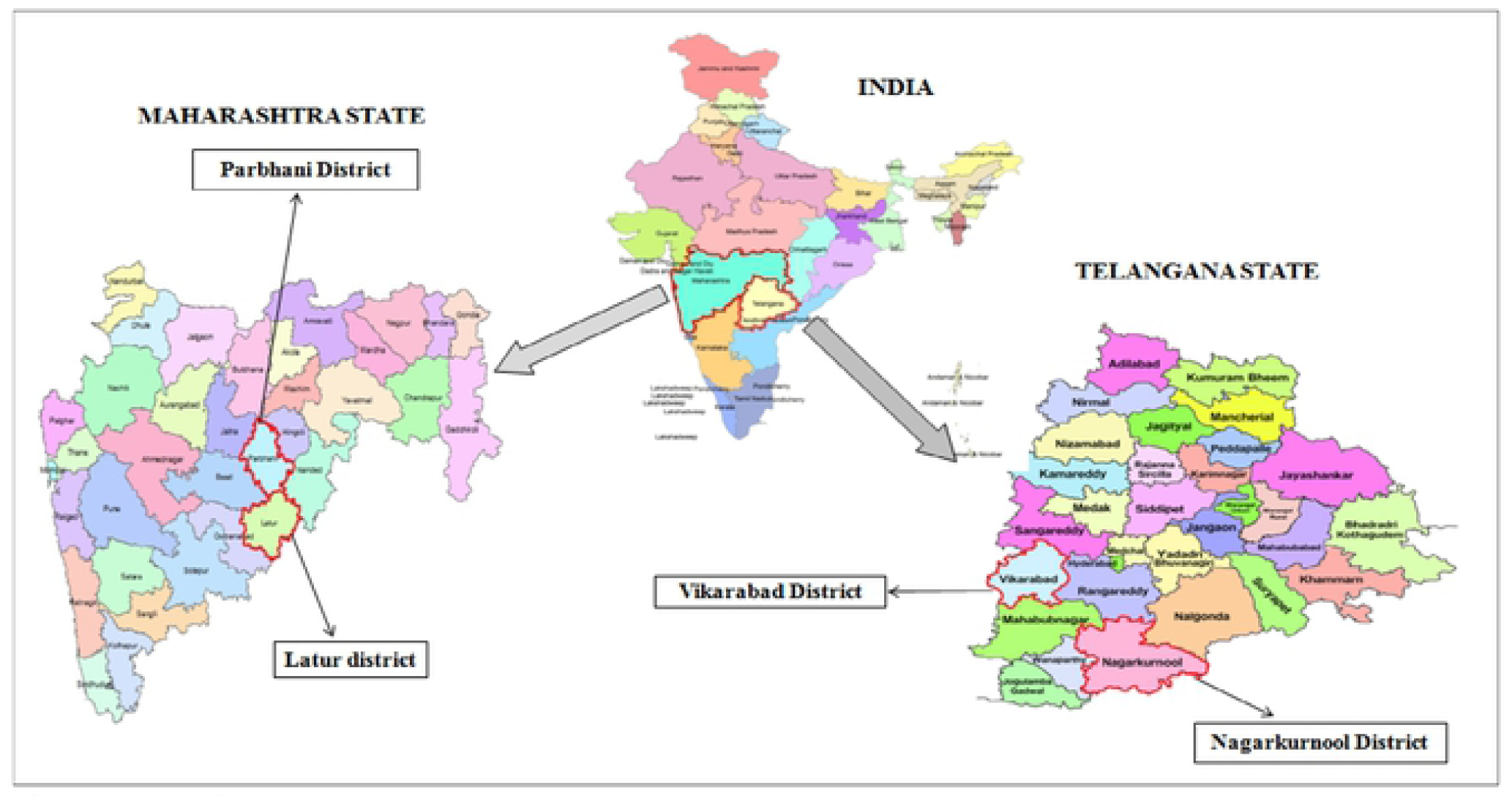
Study area.

### Profile of study zone

The study sites are located in the semi-arid region of the country and characterized by very erratic bimodal rainfall system. The selected districts of Maharashtra and Telangana are largely rainfed with average rainfall ranging between 350-1000 mm. Though irrigation sources mainly constitute canal networks, check dams, wells and bore wells, availability of water in these water points largely depends upon the amount of rainfall as well as its distribution. Pre-monsoon season is dry to very dry and the monsoon season commences mainly from mid-June - July and lasts till September end. Kharif season sowing generally begins after the onset of monsoon showers in the region. Pulses are one of the major crops in all the districts covering around 30% of the gross cropped area and more than 40 % of the total pulses area is under pigeon pea crop. The soil includes red sandy loams and red loams with clay base. Pigeon pea is the major pulses cultivated in these districts followed by green gram, Bengal gram and black gram.

### Methodology

Conjoint analysis is a popular technique used in market research with the idea that a product can be described by its attributes and their levels. In recent years, it has been applied to a number of studies to estimate the farmers’ preference towards varietal traits, seed sector, conservation incentives (Patil et al., 2006; Mafuru et al, 2007; Basavaraj et al., 2015; Conner et al., 2016,). The conjoint measurement allows estimation of the impact of individual attribute levels on the overall utility of a product (Annunziata & Vecchio, 2013)

The first step of the analysis is to identify the appropriate attributes and their levels important to the farmers. For the present study, the attributes and their levels were selected on the basis of thorough discussion with the key experts like plant breeders, agronomists and economists. Subsequently, the attributes were also discussed with pigeon pea growers in focus group discussions to validate their relevancy from the point of view of farmers. Based on the discussions, only four attributes viz. resistance (drought/ waterlogging and pod borer), yield (more than 1.5 t/ha and up to 1.5 t/ha), crop duration (short, long and medium) and plant height (short and long) were included in the study.

Out of all the possible combinations that can be made using the traits and their respective levels, eight combinations were generated using orthogonal design using SPSS software which are presented in Table 1. These combinations were presented to farmers on separate cards and they were asked to rank them on the scale of 1 to 8 (1= “most preferred” and 8 = “least preferred”). The preferences revealed by the respondent were used to estimate his part-worth utilities. The relative importance of the product attributes was estimated from part-worth utilities. It can be defined as follows:

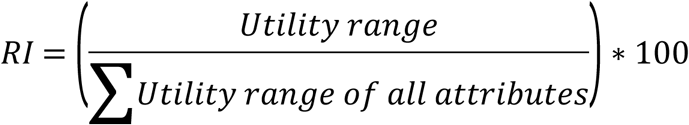

where Utility Range = Highest minus lowest utility for the attribute

**Table 1.** Attributes and their levels used for pigeon pea in conjoint analysis.

| ID assigned | Resistance | Yield | Crop duration | Plant height |
| --- | --- | --- | --- | --- |
| Card ID 1 | Drought and pod borer | More than 1.5 t/ha | Up to 140 days<br>(Short) | Up to 4 feet<br>(Short) |
| Card ID 2 | Water logging and pod borer | More than 1.5 t/ha | More than 180 days<br>(Long) | More than 4 feet<br>(Long) |
| Card ID 3 | Water logging and pod borer | Upto 1.5 t/ha | Upto 140 days<br>(Short) | More than 4 feet<br>(Long) |
| Card ID 4 | Water logging and pod borer | More than 1.5 t/ha | Upto 140 days<br>(Short) | Upto 4 feet<br>(Short) |
| Card ID 5 | Drought and pod borer | More than 1.5 t/ha | 140-180 days<br>(Medium) | More than 4 feet<br>(Long) |
| Card ID 6 | Drought and pod borer | Upto 1.5 t/ha | More than 180 days<br>(Long) | More than 4 feet<br>(Long) |
| Card ID 7 | Water logging and pod borer | Upto 1.5 t/ha | 140-180 days<br>(Medium) | Upto 4 feet<br>(Short) |
| Card ID 8 | Drought and pod borer | Upto 1.5 t/ha | More than 180 days<br>(Long) | Upto 4 feet<br>(Short) |

## Results and discussion

### Socio-economic profile of sampled farmers

The socio-economic profile of sampled farmers is presented in Table 2. The average age of the household head was 45 years and 51 years, respectively in Telangana and Maharashtra. Education status of sampled farmers of Telangana showed that 42 per cent of the sampled farmers are illiterate, while 29 per cent and 13 per cent of the farmers studied up to high school and intermediate, respectively. There were around 16 per cent of the sampled farmers who had a graduation degree. On the other hand, the percentage of illiterate farmers were lower in Maharashtra. Around 25 per cent and 43 per cent attained education up to high school and intermediate level, respectively. Whereas, 26.4 per cent of the sampled farmers attained graduation or above degree.

**Table 2.**
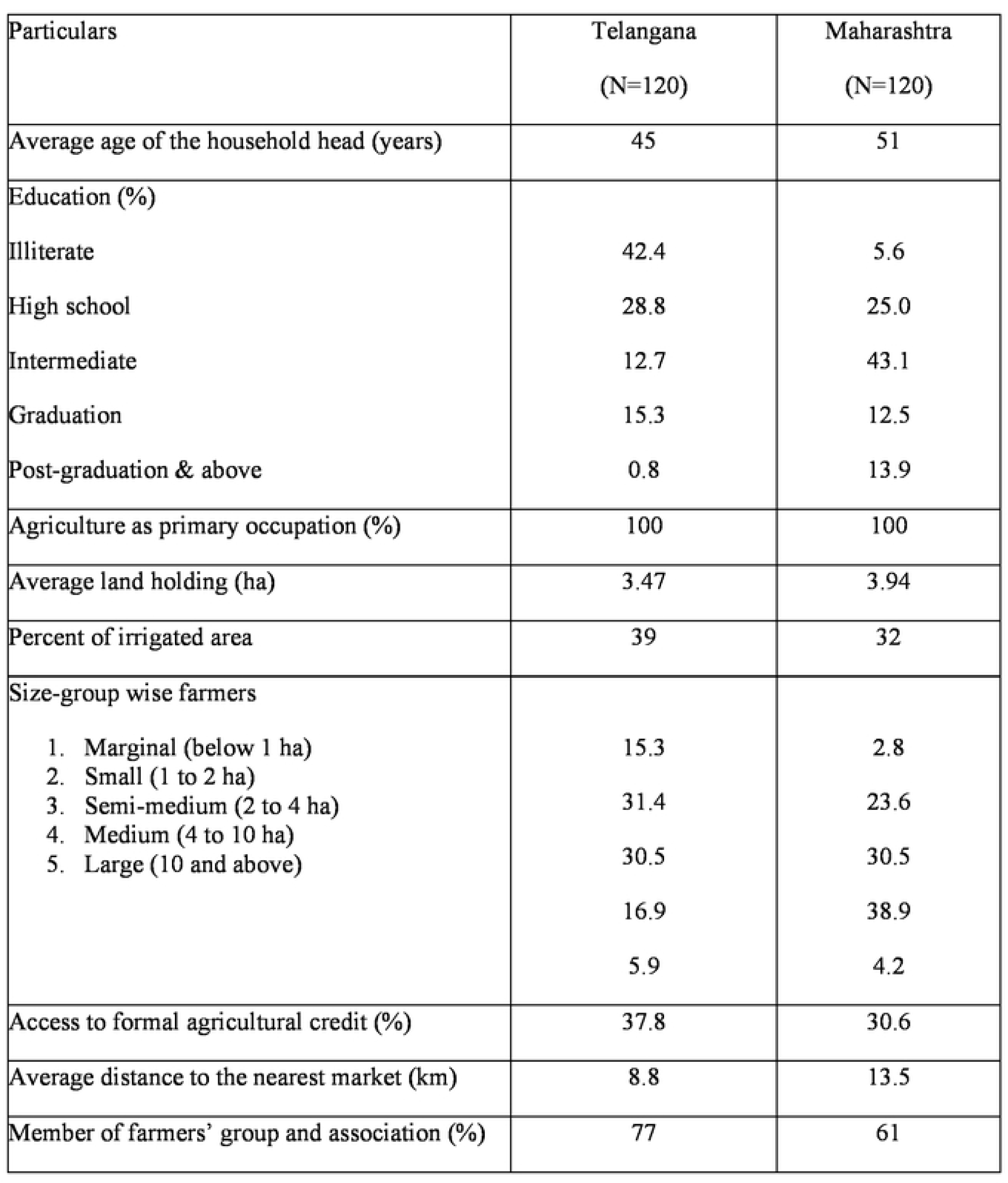
Summary characteristics of sampled pigeon pea farmers in the study area.

The average landholding size of the sampled household was found to be 3.47 ha in Telangana and 3.94 ha in Maharashtra. Only 32 and 39 percent of the holdings were reported under irrigation in Maharashtra and Telangana, respectively. Table 2 shows that average market distance of sampled growers was approximately 8-13 km from the main market in both the states and they had received at least 1 farm visit by extension officers during the cropping season. On average, 77 and 61 per cent of the sampled farmers were members of farmer groups and associations in both the states, respectively.

### Pigeon pea production constraints and coping mechanisms in the study area

Cotton, soybean, Kharif jowar, sugarcane, pulses especially pigeon pea and black gram, wheat, canola, rabi jowar and vegetables are the important crops of the region. Since most of the land is rainfed, pulses constitute a prominent position in the gross cropped area of the selected districts. Among pulses, pigeon pea suits very well to the rainfed conditions as it has deep rooting system to withstand moisture stress. Since pigeon pea is a long duration crop (almost like perennials) and is also a slow-growing crop (first four months), it is very well suited as an intercrop with any of the fast-growing annual crops like soybean, cotton, sorghum and green gram/black gram. In surveyed districts of Maharashtra, most farmers (84%) grew pigeon pea as intercrop with soybean and cotton. (Table 3).

**Table 3.** Number of farmers having pigeon pea and its combinations.

| State | Sole/Crop combination | Farmers (%) |
| --- | --- | --- |
| Maharashtra | Pigeon pea (sole) | 14 |
|  | Pigeon pea + Soybean | 64 |
|  | Pigeon pea + Cotton | 20 |
|  | Pigeon pea + Sorghum | 2 |
| Telangana | Pigeon pea (sole) | 83 |
|  | Pigeon pea + Cotton | 8 |
|  | Pigeon pea + Maize | 5 |
|  | Pigeon pea + Green gram | 4 |

In the survey, farmers were asked to rank each of the major production constraints in terms of its frequency and the impact as “most important”, “less important” and “not at all important”. Around 93 per cent farmers in Telangana and 75 per cent farmers in Maharashtra reported low rainfall or drought condition during sowing time as the most important environmental constraint in pigeon pea production (Table 4). Farmers reported heavy yield losses due to dry spell. The crop is normally sown after the onset of monsoon (second fortnight of June to the end of July) in all the districts to ensure moisture availability during germination. The majority of farmers in Maharashtra planted pigeon pea as intercrop mainly as risk minimization and crop diversification strategy. Apart from drought, few farmers in the study area reported waterlogging as another abiotic constraint affecting production adversely. Since pigeon pea is generally grown in undulated land, the waterlogging condition occurs in lowlands due to heavy rainfall.

**Table 4.**
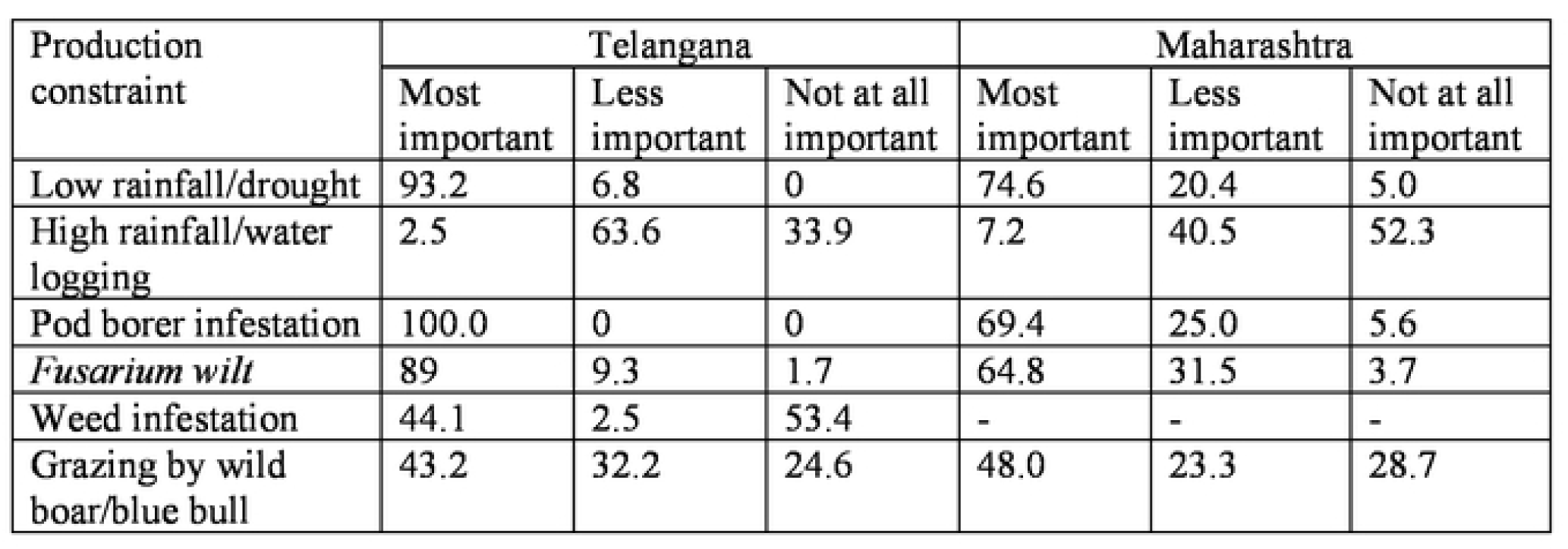
Ranking of major production constraints by the respondents(% sample households)

| Production constraint | Telangana |  |  | Maharashtra |  |  |
| --- | --- | --- | --- | --- | --- | --- |
|  | Most important | Less important | Not at all important | Most important | Less important | Not at all important |
| Low rainfall/drought | 93.2 | 6.8 | 0 | 74.6 | 20.4 | 5.0 |
| High rainfall/water logging | 2.5 | 63.6 | 33.9 | 7.2 | 40.5 | 52.3 |
| Pod borer infestation | 100.0 | 0 | 0 | 69.4 | 25.0 | 5.6 |
| *Fusarium wilt* | 89 | 9.3 | 1.7 | 64.8 | 31.5 | 3.7 |
| Weed infestation | 44.1 | 2.5 | 53.4 | - | - | - |
| Grazing by wild boar/blue bull | 43.2 | 32.2 | 24.6 | 48.0 | 23.3 | 28.7 |

The major pigeon pea pests, in decreasing order of importance, are pod borer (*Helicoverpa armigera, Maruca vitrata*), pod fly (*Melanagromyza* spp.) and white grub. Almost all the farmers in Telangana reported the yield losses due to pod borer whereas, 69.4 per cent of Maharashtra farmers reported its infestation as most important during the study period. A sizeable number of respondents reported drying up of the whole plant caused by *Fusarium wilt*, a soil-borne disease, in pigeon pea fields as the most important problem. Around 89 per cent and 65 per cent of the farmers reported the incidence of wilt in Telangana and Maharashtra, respectively. The problem of weed in pigeon pea is not very common. The weed infestation (*Cuscuta* spp) was observed only in the fields of Telangana state.

### Varietal diversity in pigeon pea

In Telangana, more than one-third of the farmers were cultivating local varieties of pigeon pea (Table 5). It was also noted that many a times farmer does not know the names of variety as normally he does not purchase the seed from formal sources. PRG-176 is the most preferred pigeon pea variety in the study area grown by 33% of the sampled farmers followed by Asha. PRG-176 was released by RARS, Palem in 2015 and is suitable for low rainfall areas with 130-135 days’ maturity. Asha is long-duration variety (180 days of maturity) released from ICRISAT in 1993. Only 6 per cent of the sampled farmers were growing Durga variety of pigeon pea. This variety escapes from the pod borer damage as it flowers and bears pods before the dreaded pest peak.

**Table 5.** Prominent varieties cultivated by the sampled farmers.

| State | Pigeon pea variety grown | Year of release | Farmers growing the crop variety (%) | Source of seed (%) |  |
| --- | --- | --- | --- | --- | --- |
|  |  |  |  | Own/home saved | Purchased |
| Telangana | Local | - | 35 | 76 | 24 |
|  | PRG-176 | 2015 | 33 | 40 | 60 |
|  | Asha (ICPL 87119) | 1993 | 20 | 76 | 34 |
|  | Durga (ICPL 84031) | 1995 | 6 | 39 | 61 |
|  | PRG-158 | 2007 | 4 | 0 | 100 |
|  | Laxmi (ICPL 85063) | 2002 | 2 | 100 | 0 |
| Maharashtra | Local (Nirmal, Yashodha) | - | 59 | 47 | 53 |
|  | BSMR 736 | 1996 | 23 | 72 | 28 |
|  | Maruti (ICPL-8863) | 1986 | 10 | 67 | 33 |
|  | BSMR 853 | 2002 | 5 | 63 | 37 |
|  | Others (BDN 711, BDN 2) | - | 3 | 55 | 45 |

In Maharashtra, 59 per cent of the farmers were cultivating local varieties viz. Nirmal and Yashodha. These varieties are being marketed by private seed companies and are mainly preferred due to their high yielding and early maturity attributes. The early maturity trait is extremely important as it helps the crop to escape from terminal water stress. Their harvesting starts by the end of November. Apart from local varieties, BSMR-736 is the most preferred variety followed by Maruti. Varieties like BSMR 853 (white seeded), BDN 711 and BDN 2 were also being grown in the area. It was found in the study that majority of the farmers were not purchasing seeds from the market instead of using own home saved seeds or exchanging with fellow farmers, which also reflects imperfection in the seed market. The other important sources of seed purchases were government agencies and private seed dealers.

### Pigeon pea varietal traits preferences of the sample farmers

Conjoint analysis is used to understand what specific values of attributes farmers considers while deciding regarding the cultivation of a crop variety. The farmers were asked to reveal their preferences for various attributes of pigeon pea in the form of ranks to eight combinations. The utility derived by the farmer from the attributes are estimated (Table 8) and state-wise relative importance of different attributes is depicted in Figure 2. Perusal of the figure reveals that farmers’ perception and ranking of the different attributes varied across the study sites. Farmers of Telangana assigned the highest importance to crop duration followed by resistance to pod borer and drought, yield and the plant height respectively. On the other hand, sample respondents of Maharashtra state preferred resistance to drought and pod borer trait the most followed by crop duration, yield and plant height. This suggests that the farmers’ preferences lie more with pigeon pea yield and its stability attribute in both the regions. The least important attribute in pigeon pea variety as per farmers’ perception is plant height.

**Figure 2:**
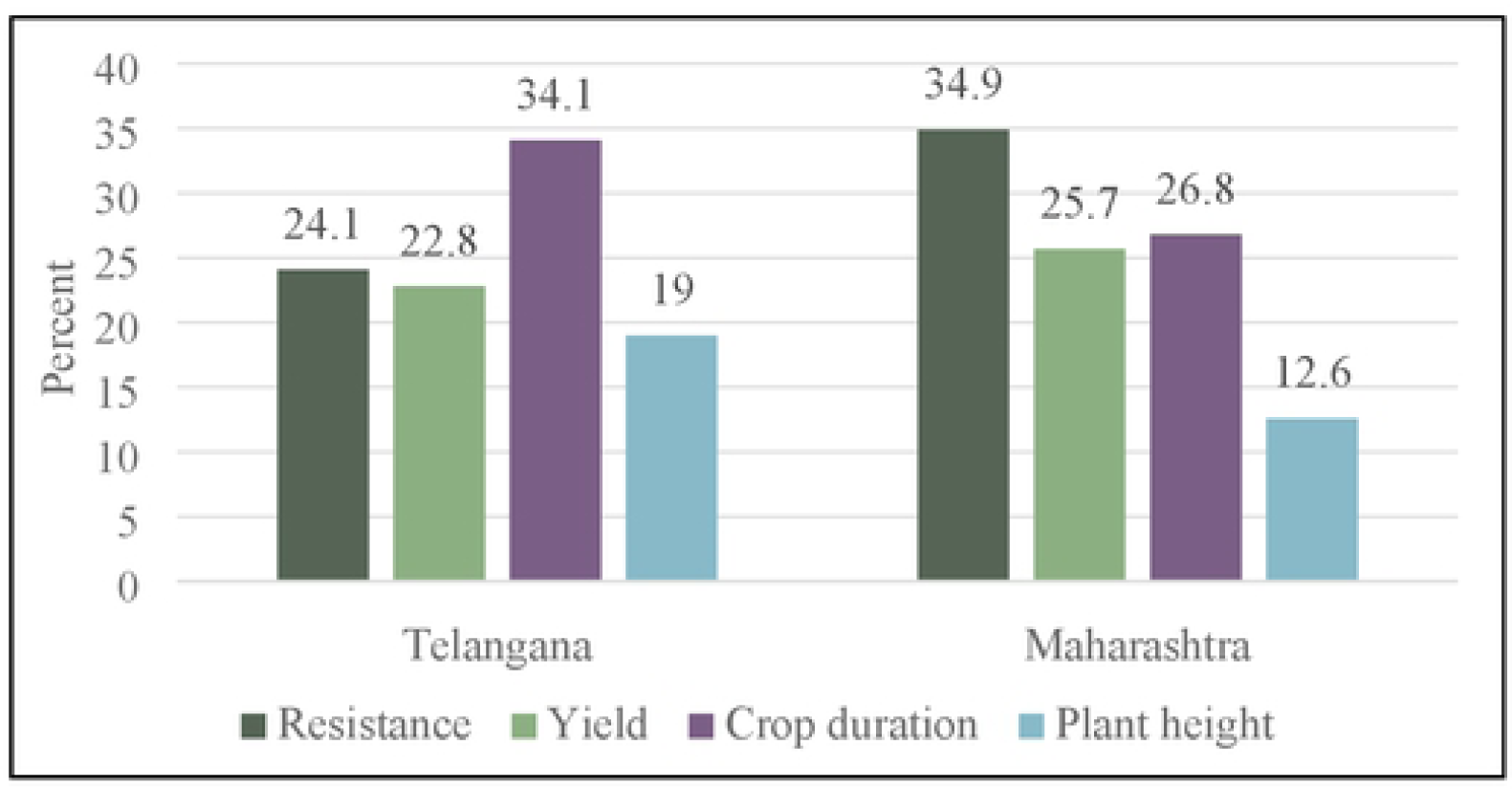
State-wise relative importance of pigeon pea attributes given by farmers.

The part-worth estimates for different levels of pigeon pea attributes are given in Table 6. The negative part-worth indicates the negative preference of the farmers towards that level of the attribute. The value attached to part-worth represent the magnitude of utility derived from that specific level. In crop duration, short-duration variety is preferred by the farmers and its contribution to the total utility is 0.390 and 0.374 in Telangana and Maharashtra, respectively. The short duration varieties would help the farmer avoid losses from terminal stress during winters in the crop as well as fetching higher market prices in the mandi due to early arrival. Farmers prefer pigeon pea variety with yield more than 1.5 t/ha. Since the study area is largely rainfed, farmers prefer variety with drought and pod borer resistance. The farmers of both states preferred long

**Table 6.**
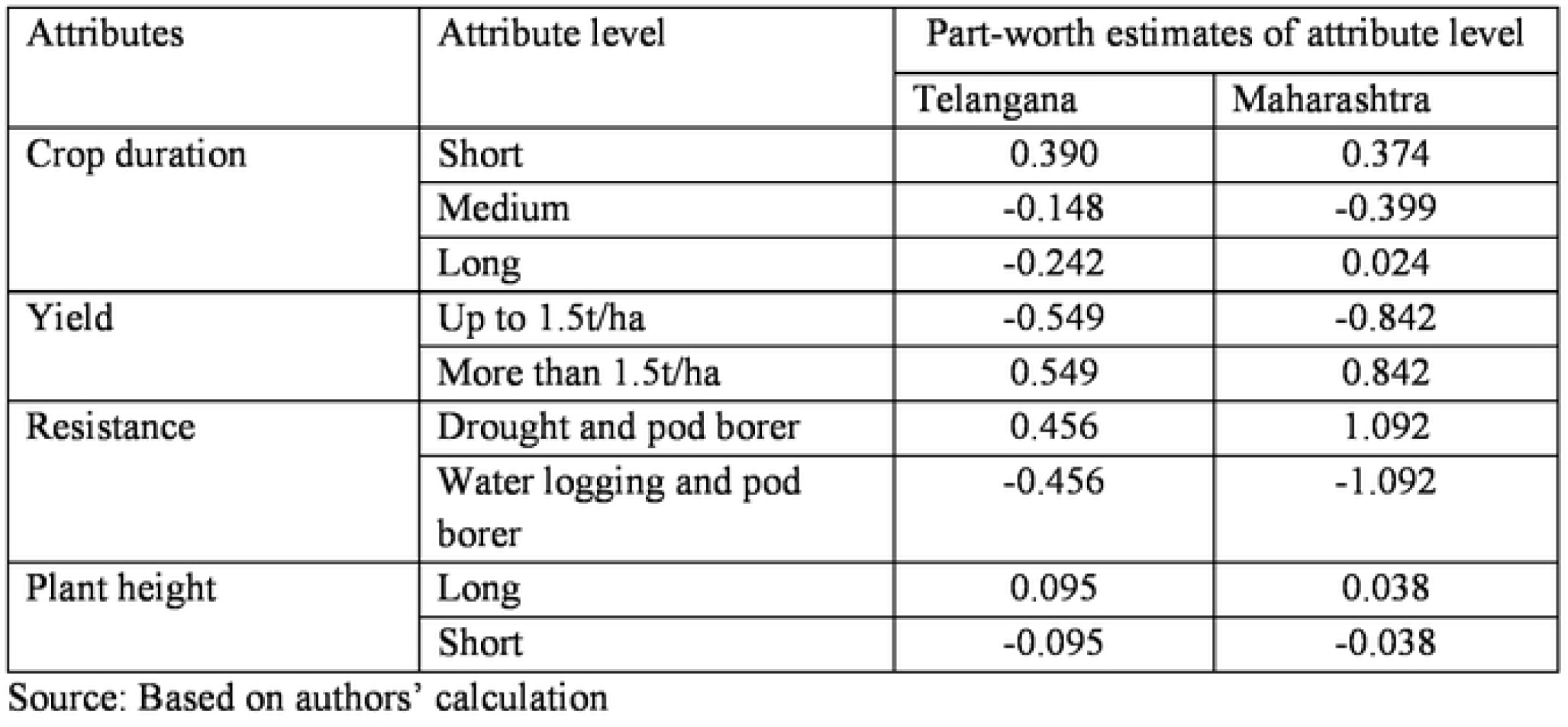
Part-worth estimates of attributes of pigeon pea variety for the farmers.

| Attributes | Attribute level | Part-worth estimates of attribute level |  |
| --- | --- | --- | --- |
|  |  | Telangana | Maharashtra |
| Crop duration | Short | 0.390 | 0.374 |
|  | Medium | -0.148 | -0.399 |
|  | Long | -0.242 | 0.024 |
| Yield | Up to 1.5t/ha | -0.549 | -0.842 |
|  | More than 1.5t/ha | 0.549 | 0.842 |
| Resistance | Drought and pod borer | 0.456 | 1.092 |
|  | Water logging and pod borer | -0.456 | -1.092 |
| Plant height | Long | 0.095 | 0.038 |
|  | Short | -0.095 | -0.038 |

Source: Based on authors' calculation

### Processors’ and consumers’ preferences of pigeon pea varietal traits

India is the largest producer and consumer of pulses and dhal mills are major constituent of the food processing industry. Although production is scattered across the country, the organized dhal processing industry is located in selected pockets of the country. According to India Pulses and Grains Association (IPGA), about 75 per cent of pulses are processed by organized Dhal mills, which increased from 2000 dhal mills in 1972 to 14000 mills in recent years. Dhal mills hubs are located in pulses production centres such as Indore, Jalgaon, Akola and Nagpur as well as near major consumptions centres like Kolkata, Chennai, Mumbai, Hyderabad and Delhi. Maximum number of dhal mills are located in the state of Maharashtra (major pigeon pea producing state) with an average milling capacity of 18 tons per day. Chennai follows next and home to 750 dal mills followed by Delhi with 300 dal mills. Besides these, the traditional dal mill or home-based manual processing also plays an important role in dal processing.

The preferred varietal traits were ranked by the processor/dal millers located in Nagpur, Akola, and Latur districts in Maharashtra. The nature of the seed coat was the most preferred character of the variety as it determines the duration and cost of the processing. It is to be noted that among the pulses, the duration of the pigeon pea processing is very lengthy (about 8-9 days), whereas in case of chickpea it is just 2-3 days. Among varieties, Maruti (ICP 8863) is still the preferred variety by the processors as it is the best suited for processing. Dal recovery percentage is the major factor which influences the purchase behaviour of the processor. As of now, 70-72 per cent recovery is made by the processors and remaining are powder and husk in equal share. These by-products are preferred mainly for cattle feed and fetch less price. Processors’ demand more R&D on by-products to increase their market value and food chain of these products. Third, uniformity of the grain size is very important attributes for processing as heterogeneous grains lot produce more unprocessed seeds resulting in higher processing cost. As far as size is concerned, medium-sized grains are preferred when compared to the large or small grains as the consumers demand medium-sized dal for consumption.

Consumers, on the other hand, find traits such as clean, uniform grains and less cooking time more desirable. Pigeon pea is consumed regularly, over half of the respondents eat it daily or several times a week. In the household, women (95%) decided which pulses to eat and how to prepare it. 96% of the respondents use pigeon pea to prepare dhal. Their awareness about other traits which have directly impacted pigeon pea productivity in India was minimal, 87% of consumers did not know any pigeon pea varieties.

## Conclusion

This study assesses India’s food security implications due to stagnating and low yield of pigeon pea and ascertain farmers’ preferences of pigeon pea varietal traits, production constraints and farmers’ coping strategies in diverse pigeon pea production environments. The study also aimed at identifying market preferred seed types and traits through detailed survey of processors and consumers. The study determined the types of pigeon pea varieties farmers grow, farmers’ and processors’ preferences of the grain type, varieties & constraints to production, on which basis research strategies for improvement of pigeon pea production could be formulated. Intercropping of medium to long duration pigeon pea varieties with soybean, cotton and sorghum and sole cropping of landraces and improved varieties are the dominant production system in central and southern zones of the pigeon pea growing areas. However, the results also indicated very low varietal diversity and prevalence of old varieties in all the regions.

Production constraints in these two three regions were basically similar, with most of the farmers identifying pod borer & pod fly as the major pest, and wilt as the major disease. Drought was identified by almost all the farmers as a major production constraint. Farmers indicated the use of clean seed, high yielding varieties, inter & mixed cropping, planting density, and manure application as some of the strategies they used to cope with the production constraints. In terms of preference for new cultivars, farmers preferred high grain yield with drought tolerance, medium to early maturity, pod borer resistance, tolerance to wilt disease, moderate plant height and ease of threshing without compromising other preferred attributes. The study also revealed that a considerable proportion of the farmers had limited knowledge of improved varieties, control strategies for pod borer and wilt disease. This therefore calls for more research and adequate extension services for better farm management. The analysis of processors’ preference of grain types for dal processing showed that uniform size, oval shape, orange coloured seed and most importantly moderately hard seed coat are highly desirable. Consumer preference is more towards uniform seed size and less cooking time.

## Acknowledgement

The authors gratefully acknowledge funding for this study from the Indo-Swiss Collaboration in Biotechnology (ISCB).

## Footnotes

1 Dhal is made by removing the seed coat and splitting the cotyledons. Dhal made from pigeon pea is referred to as *Tur dhal*, although the terms *Toor*, Tuver or *Arhar dhal* are also used to describe the same product.

